# Electrically Controlling and Optically Observing the Membrane Potential of Supported Lipid Bilayers

**DOI:** 10.1101/2021.12.10.472087

**Authors:** Shimon Yudovich, Adan Marzouqe, Joseph Kantorovitsch, Eti Teblum, Tao Chen, Jörg Enderlein, Evan W. Miller, Shimon Weiss

## Abstract

Supported lipid bilayers are a well-developed model system for the study of membranes and their associated proteins, such as membrane channels, enzymes, and receptors. These versatile model membranes can be made from various components, ranging from simple synthetic phospholipids to complex mixtures of constituents, mimicking the cell membrane with its relevant physiochemical and molecular phenomena. In addition, the high stability of supported lipid bilayers allows for their study via a wide array of experimental probes. In this work, we describe a platform for supported lipid bilayers that is accessible both electrically and optically. We show that the polarization of the supported membrane can be electrically controlled and optically probed using voltage-sensitive dyes. Membrane polarization dynamics is understood through electrochemical impedance spectroscopy and the analysis of the equivalent electrical circuit. We also describe the effect of the conducting electrode layer on the fluorescence of the optical probe through metal-induced energy transfer. We conclude with a discussion on possible applications of this platform for the study of voltage-dependent membrane proteins and other processes in membrane biology and surface science.

## Introduction

Biological membranes are a main component of life, providing cells and membrane-enclosed organelles compartmentalization, together with functional, communicational, and computational capabilities. By utilizing various phospholipids, glycolipids, cholesterol and proteins, complex lipid bilayers acquire different physiochemical and molecular properties that allow for intricate functions, such as chemical and electrical gating, signal transduction, and catalysis. Among several model lipid bilayer systems, supported lipid bilayers (SLBs) [1, 2] offer an extremely versatile and robust platform for the study and application of membranes and associated lipids and proteins [3–5].

At its simplest form, SLB systems are composed of phospholipids deposited on top of a solid support to make a continuous planar bilayer. Planar bilayer formation can be achieved by, for example, the Langmuir-Blodgett technique [6, 7], the adsorption and fusion of vesicles [8] or bicelles [9], or by gradual organic to water solvent exchange [10]. The physiochemical properties of the supported bilayer, such as its phase and fluidity, will be determined by the composition of the bilayer [11] and the properties of the underlying substrate [12]. These properties, in turn, can influence the structure and functionality of embedded proteins. In addition, nuclear magnetic resonance [13], neutron [14] and x-ray reflectometry [15] showed that a hydration layer of 5-30 Å in thickness is present between the substrate and bilayer, influencing the fluidity of the lower leaflet. The artificial interface occurring at the leaflet closer to the support can prevent membrane proteins with extracellular domains to be fully functional, as they can become immobilized and denatured due to the interaction between the protruding hydrophilic domain and the nonphysiological underlying environment [3]. In order to lessen the adverse effects of the artificial solid support, hydrated polymer films can be deposited on the support prior to the bilayer formation [16, 17]. In addition to cushioning polymer films, many approaches for tethering lipid bilayers to the solid support have been developed [18]. For example, DNA-lipid conjugates can be used in order to distance the bilayer from the solid support in a controllable manner [19]. Due to the high stability of SLBs, many experimental techniques can be applied in order to study the membrane and its associated biomolecules. In particular, the electrochemical properties of SLBs were thoroughly studied using different supporting substrates, such as doped silicon [20, 21], silicon-silicon dioxide [22], indium tin oxide (ITO) [21, 23–25], titanium-titanium dioxide [26], gold [21], gold-silicon dioxide [27, 28], and semiconducting polymers [29].

The electrical potential across a lipid bilayer dictates many of its functional properties and is usually defined in three main regions [30–32]: 1. The potential arising from the charged headgroups and adsorbed ions at the membrane-liquid interface (surface potential). 2. The potential produced by the aligned molecular dipoles of both lipids and water at the interface (dipole potential). 3. The overall potential difference due to the different ion concentrations on both sides of the membrane (transmembrane potential). Transmembrane potential plays a main role in the communicational and computational capabilities of cells through its effect on membrane proteins. An important and highly studied class of membrane proteins is voltage-gated ion channels. These transmembrane proteins contain voltage-sensing gate domains that translocate due to applied electric fields, thus regulating ion conductance [33, 34]. Beyond voltage-gated ion channels, numerous transporters [35] and enzymes [36, 37] are directly affected by the transmembrane potential. In addition, it is very likely that many proteins and processes in the membrane are affected or induced by the electrical potential across the membrane [38, 39]. In particular, microscopic electrostatics can direct the conformation and function of membrane proteins through their charged domains [40, 41].

In this work, we describe a platform for SLBs that is addressable both electrically and optically. We characterize the electrical response of the bilayer with electrochemical impedance spectroscopy (EIS) [42] and demonstrate the ability to polarize the membrane electrically while optically monitoring the membrane potential using voltage-sensitive membrane dyes. We show that the optically measured changes in membrane potential can be understood through the dynamical properties of the model equivalent electric circuit. In addition, we describe the effect of the conducting electrode layer on the radiative transition of the demonstrated membrane dye through metal-induced energy transfer (MIET) [43, 44], and discuss its implications. Finally, we present possible future applications for this platform, emphasizing the complementarity between optical and electrochemical techniques and their relevance in the study of voltage-dependent membrane proteins and other voltage-dependent processes and phenomena in molecular biology and surface science.

## Materials and methods

### Metal-Oxide Substrate

Glass coverslips (Marienfeld Superior, Lauda-Königshofen, Germany) coated with a layer of ~8 nm Ti and ~20 nm Ta_2_O_5_ served as an optically transmissive metal-oxide substrate for the lipid bilayer for all the experiments described in this work, apart from Figs. 5 and S4, where varying thicknesses of Ta_2_O_5_ were used. Deposition of both Ti and Ta_2_O_5_ layers was performed using ion beam sputtering technique (Nanoquest I; Intlvac, Ontario, Canada). For the Ti layer, a stainless steel mask (3 mm in diameter) was used in order to define the surface area of the electrode. Ti was sputtered from a Ti target at an average sputtering rate of ~1 Å/s, using a sputtering ion beam voltage of 1200 V and beam current of 120 mA, with Ar (10 sccm) as the working gas. Ta_2_O_5_ was sputtered from a Ta target at an average sputtering rate of ~0.9 Å/s, using a sputtering ion beam voltage of 1200 V and beam current of 110 mA, with Ar (10 sccm) as the working gas, and an assisting ion source composed of Ar (10 sccm) and O_2_ (18 sccm) at an ion beam voltage of 120 V and beam current of 95 mA.

Substrate roughness characterization was performed in air with atomic force microscopy (AFM). AFM measurements were carried out using a scanning probe microscope (Bio FastScan; Bruker, Karlsruhe, Germany). All images were obtained using soft tapping mode with a silicon probe (Fast Scan B; Bruker) with a spring constant of 1.8 N/m. The resonance frequency of the cantilever was approximately 450 kHz (in air). The images were captured in the retrace direction with a scan rate of 1.6 Hz, with a resolution of 512 samples per line. Image processing and roughness analysis were performed using a dedicated open-source software (Gwyddion) [45].

### Lipid Vesicle Preparation

Homogenous lipid vesicles were prepared by hydration of a dry lipid mixture, followed by sonication and extrusion. In short, 530 nmol of 1-palmitoyl-2-oleoyl-glycero-3-phosphocholine (POPC; Avanti Polar Lipids, Alabaster, AL) and 150 nmol of 1-palmitoyl-2-oleoyl-sn-glycero-3-phospho-L-serine (POPS; Avanti Polar Lipids, Alabaster, AL) were dissolved in chloroform and mixed in a molar proportion of approximately 80% and 20%, respectively, evaporated for 1 h at 40°C using a centrifugal vacuum concentrator (CentriVap Concentrator; Labconco, Kansas City, MO), and dried under vacuum overnight. The lipid film was dispersed in 1 ml of Tris-NaCl buffer (10 mM Tris, 100 mM NaCl, pH 8), giving a final lipid concentration of ~ 0.6 mM. The solutions were sonicated by a probe sonicator for 30 seconds at 26 Watt, 20 kHz (VCX 130; Sonics, Newtown, CT). Afterward, the lipid solutions were centrifuged for 20 min at 16,000 x g and 4 °C. The suspensions were then extruded (Mini-Extruder; Avanti Polar Lipids, Alabaster, AL) 11 times through 100 nm polycarbonate membranes (Nuclepore Hydrophilic Membrane; Whatman, Clifton, NJ). The hydrodynamic size distribution of the vesicles at each step was measured using dynamic light scattering (Zetasizer Nano ZS; Malvern Instruments, Westborough, MA) using a 633 nm laser and a scattering angle of 173°, operating at 25 °C (Fig. S1). In order to ensure repeatability between preparations, the total scattered intensity was used as an indicator for the vesicle concentration.

### Supported Lipid Bilayer Formation

A POPC:POPS lipid bilayer with an overall negative surface charge was prepared by the rupturing and fusing of lipid vesicle on the hydrophilic metal-oxide substrate. Substrates were first cleaned by 5 min sonication in detergent solution (2% Hellmanex III; Hellma, Müllheim, Germany), double-distilled water (DDW), and acetone. Next, the substrates were rinsed with DDW and dried under a stream of N_2_. In order to ensure a high degree of hydrophilicity, substrates were further treated under a glow discharge system (Emtech K100x; Quorum Technologies, Sussex, UK) for a total of 5 minutes. Next, a disposable polylactic acid (PLA) chamber (shown in Fig. 1A) was attached on top of the treated substrate using a silicone sealant glue. The experimental chamber was immediately transferred to a vacuum desiccator for 20 min, allowing the sealing agent to dry while keeping the treated electrode surface unaffected by humidity. Next, 50 µL of the vesicle solution was added to the chamber, together with Tris-NaCl buffer containing CaCl_2_ for a final concentration of 2 mM CaCl_2_. The vesicles were allowed to incubate for 20 min, followed by multiple buffer exchanges for the removal of unbound vesicles. While membrane resistance *R*_*m*_ was found to be variable between preparations, the likelihood of achieving high *R*_*m*_ considerably increased for membranes that were allowed to stabilize at room temperature for 16-24 hours after the removal of unbound vesicles. SLBs were stained with VF2.0.Cl or VF2.1.Cl shortly before experiments by incubating the SLB for 5 min in a buffer solution containing the membrane dye at a final concentration of 100 nM. Following incubation with the membrane dye, multiple buffer exchanges were performed in order to remove free dye from the buffer solution.

**Fig. 1.**
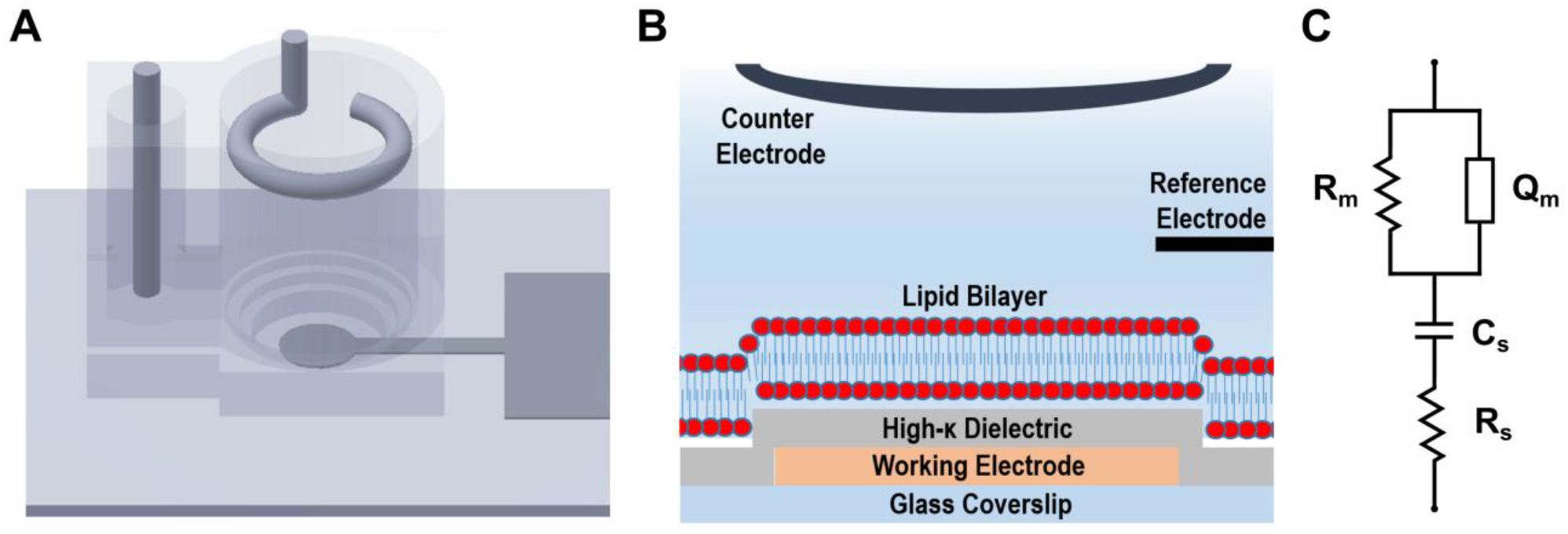
Electrically and optically addressable solid-supported lipid bilayer. **(A)** An electrochemical chamber is placed on top of a semi-transparent Ti/Ta_2_O_5_ electrode that supports the membrane. The chamber is filled with an electrolyte solution, and counter (Pt) and reference (Ag/AgCl) electrodes are positioned in the solution, forming a three-electrode cell. **(B)** A schematic drawing (not to scale) of the membrane and the underlying electrode support. **(C)** The electrical equivalent circuit model. The substrate is represented by the resistance of the semi-transparent conducting layer and the electrolyte solution *R*_*s*_ and the capacitance of the insulating layer *C*_*s*_. The membrane is approximated by a resistive element *R*_*m*_ connected in parallel to a constant phase element *Q*_*m*_.

### Electrochemical Impedance Spectroscopy (EIS)

Electrochemical characterization and control were performed in a three-electrode cell configuration using a commercial potentiostat (Autolab PGSTAT302N; Metrohm, Utrecht, Netherlands), with the conductive Ti substrate connected as the working electrode, a Pt wire as the counter electrode, and an Ag/AgCl electrode as the reference electrode. Impedance spectroscopy was performed using a 10 mV AC modulation amplitude at 0 V bias voltage. EIS measurements of the supporting electrode, without the membrane, were performed at the end of each experiment by the removal of the supported membrane via multiple washing steps using detergent and DDW, followed by buffer exchange to Tris-NaCl buffer. All measurements were performed in 10 mM Tris with 100 mM NaCl (pH 8).

### Fluorescence intensity and lifetime measurements of membrane potential

The fluorescence intensity and lifetime of the membrane dyes VF2.0.Cl and VF2.1.Cl was measured using a time-resolved imaging system based on an inverted microscope (IX83; Olympus, Tokyo, Japan) coupled to a 60X water-immersion objective (UPLSAPO60XW; Olympus). A light-emitting diode light source (SPECTRA X; Lumencor, Beaverton, OR) and a sCMOS camera (ORCA-Flash4.0 V3; Hamamatsu, Tokyo, Japan) were used for epifluorescence imaging. Time-resolved confocal fluorescence detection was performed in a time-correlated time-tagged recording mode using a 470 nm pulsed diode laser (LDH-D-C-470; PicoQuant, Berlin, Germany). The emitted light was collected and focused using a 180 mm lens (AC508-180-A, Thorlabs, Newton, NJ, USA) into a 150 µm pinhole (~9 Airy units) at the conjugate image plane, and then collected and focused onto an avalanche photodiode detector (SPCM-AQRH-43-TR; Excelitas Technologies, Waltham, MA). The detector and the laser driver (PDL 800-D; PicoQuant) were connected to a time-correlated single-photon counting card (TimeHarp 260 PICO; PicoQuant). A data acquisition (DAQ) device (NI USB-6356; National Instruments, Austin, TX), controlled by home-written software (LabVIEW; National Instruments), was used for synchronizing the potentiostat and the time-correlated single-photon counting electronics. Lifetime curve fitting was performed using the maximum likelihood estimation method [46] with a model function of the form exp(−*t*/*τ*) for mono-exponential decay and ~exp(−*t*/*τ*_1_) + *R*_0_exp(−*t*/*τ*_2_) for bi-exponential decay, with an added constant background noise with a uniform probability distribution. Since the instrument response function of the described system was both temporally unstable and relatively long (~310 ps), the fitting procedure was performed on the decaying tail of the fluorescence lifetime curve.

## Results and discussion

### Membrane-supporting metal-insulator-electrolyte capacitor

An overall negatively charged lipid bilayer composed from a mixture of POPC and POPS phospholipids in a 4:1 molar ratio was formed on top of a Ti/Ta_2_O_5_ electrode (Fig. 1), as described in ‘Materials and methods’. Metal-oxide surfaces provide several key advantages as solid support for artificial lipid bilayers compared to purely conductive surfaces. First, faradic currents at the metal-liquid interface can induce electrochemical reactions with toxic products [47], which are significantly reduced once the conductive layer is passivated by an inert oxide layer. Second, when using fluorescent probes, an oxide layer reduces their undesired quenching due to energy transfer to surface plasmons in the metal layer [44, 48]. Ta_2_O_5_ was chosen to serve as the insulating layer due to its relatively high (above 20) dielectric constant, high breakdown threshold, low leakage current, chemical stability and biocompatibility [49, 50]. The nanoscale roughness of the solid support has a significant effect on the structural and dynamical properties of the SLB. The homogeneity of the bilayer film and the liquid-phase diffusional properties are related to the underlying substrate’s physical and chemical properties [51]. Even the surface treatment used prior to the deposition of the lipid membrane greatly affects membrane fluidity [52]. Since lipid membranes are 4-5 nm in thickness, the supporting substrate roughness is a crucial parameter [12], which must be minimized to a sub-nm scale in order to achieve a complete and homogeneous lipid bilayer film. While ITO is routinely used for semi-transparent conducting electrodes, its nanoscale roughness (>1 nm) limits the homogenous coverage of fluid lipid bilayers [23]. Following a screening of different possible conducting layers, we chose a thin layer (~8 nm) of Ti as the semi-transparent conductive layer due to its low roughness and good adhesion to glass and to the upper Ta_2_O_5_ layer. The surface roughness of the Ti/Ta_2_O_5_ electrode surface was characterized with AFM and measured to be below 0.5 nm (Fig. S2A). Since the phase transition temperatures of both POPC and POPS are below room temperature, the supported membrane is expected to be in a fluid phase, displaying translational diffusion of the constituting lipids. Therefore, fluorescence recovery after photobleaching (FRAP) was confirmed at several locations of each supported bilayer under study in order to ensure homogenous membrane preparations (Fig. S2B).

### Electrical characterization of the supported membrane

The electrical properties of the lipid bilayer were characterized using EIS, an alternating current (AC) technique that is commonly used to measure the frequency-dependent complex impedance of electrochemical systems. Although these measurements are relatively straightforward, the interpretation of impedance spectra and the equivalent model circuit are challenging due to the complicated nature of real systems and the ambiguity in their multivariable description. Importantly, one needs to ascribe a physical meaning or purpose to each electrical element of the equivalent model circuit.

The metal-insulator-electrolyte system can be approximately modeled as a serially connected ideal resistor *R*_*s*_, representing both the solution and substrate resistance, and capacitor *C*_*s*_, representing the insulating layer (Figs. 2 and S3C). We note that the electrical double layer at the oxide-electrolyte interface, commonly described as a combination of an immobile Stern layer and a mobile diffusive layer by the Gouy-Chapman-Stern model, is usually required for a complete description of electrode-electrolyte interfaces. These models require at least one *ad hoc* constant phase element (CPE). However, since the electrical double layer capacitance ranges between 6 to 12 µF cm^−2^ at the relevant electrolyte concentrations [53], higher than the capacitance of both the insulating layer and the membrane, its spectral contribution occurs at frequencies lower than the range described here. The impedance spectra of Ti*/*Ta_2_O_5_ electrodes with varying thicknesses of the insulating layer are shown in Fig. S3.

**Fig. 2.**
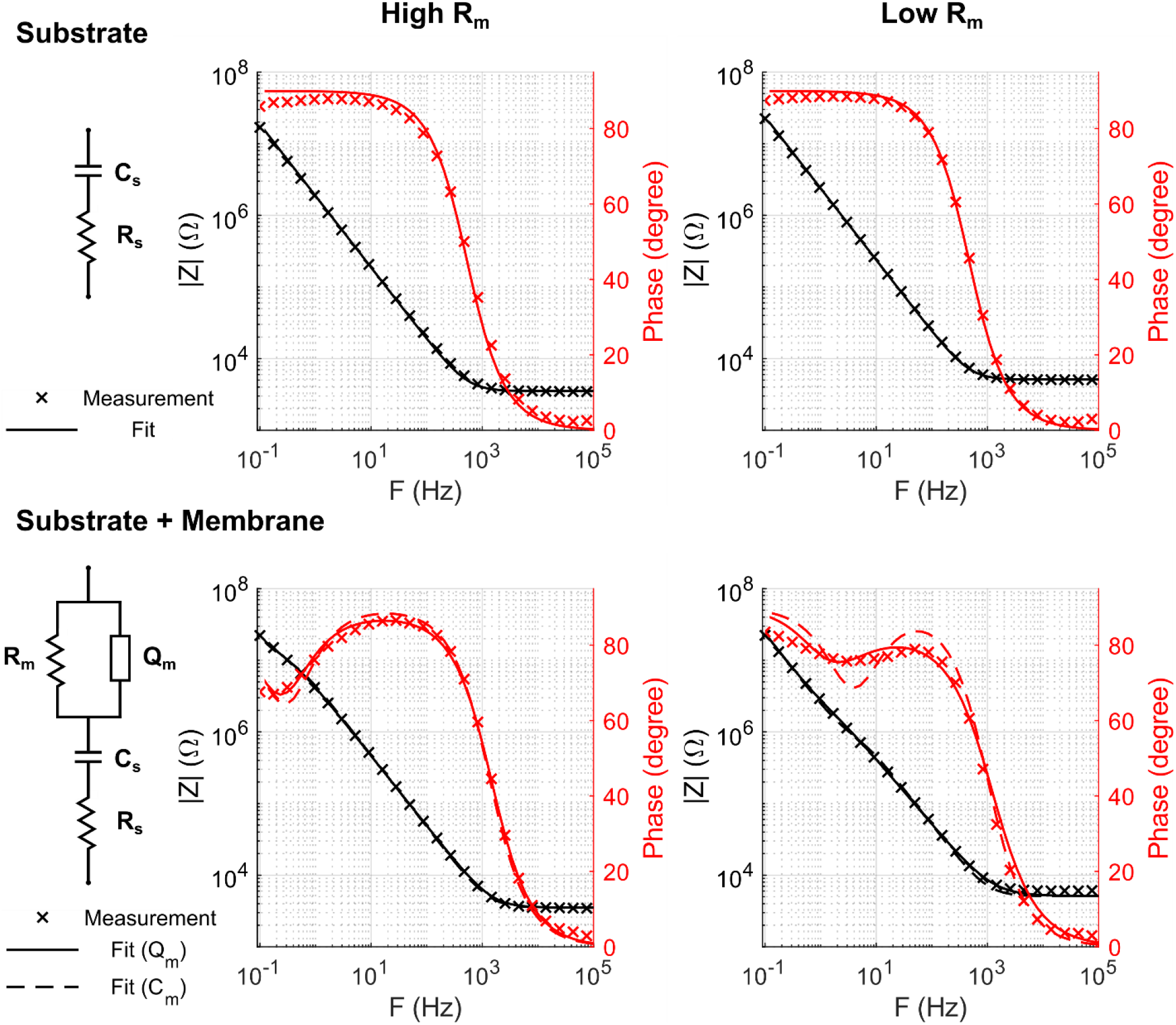
Electrical characterization of the supported membrane via electrochemical impedance spectroscopy. The complex impedance spectra (amplitude and phase) of a bare Ti/Ta_2_O_5_ substrate (top) and the same substrate supporting a lipid bilayer (bottom) are shown for two distinct membrane preparations, differentiated mainly by the time they were allowed to relax on the substrate after membrane formation, resulting in different membrane resistance *R*_*m*_. The left panel shows the equivalent circuit used in modeling the impedance spectra. The bare electrode is modeled by a simple resistive element *R*_*s*_ in series with a capacitive element *C*_*s*_, and the membrane is approximated by a resistive element *R*_*m*_ in parallel with a constant phase element *Q*_*m*_. The best fit of each impedance spectra is shown for both a general constant phase element (solid line), and the ideal case (n=1), representing a simple capacitive element 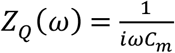 (dashed line). The fitted parameters of the bare substrate model were used as fixed parameters for the fitting of the supported membrane. The best fit parameters for the high *R*_*m*_ system are *R*_*s*_ = 249.4 Ω *cm*^2^, *C*_*s*_ = 1.2 *μ*F *cm*^−2^ for the substrate, and *R*_*m*_ = 510.5 *k*Ω *cm*^2^, *Q*_*m*_ = 0.95 *μ*F *s*^*n*−1^ *cm*^−2^, *n* = 0.96 or *R*_*m*_ = 457.9 *k*Ω *cm*^2^, *C*_*m*_ = 0.79 *μ*F *cm*^−2^ for the membrane. The best fit parameters for the low *R*_*m*_ system are *R*_*s*_ = 359.9 Ω *cm*^2^, *C*_*s*_ = 0.95 *μ*F *cm*^−2^ for the substrate, and *R*_*m*_ = 63.9 *k*Ω *cm*^2^, *Q*_*m*_ = 2.1 *μ*F *s*^*n*−1^ *cm*^−2^, *n* = 0.85 or *R*_*m*_ = 33.2 *k*Ω *cm*^2^, *C*_*m*_ = 0.84 *μ*F *cm*^−2^ for the membrane.

Several experimental studies showed that while SLBs commonly have a capacitance of 0.5 to 1 µF cm^−2^ [20, 21, 28], as would be theoretically expected for a typical layer of a lipid bilayer, membrane resistance greatly varies between different membrane compositions, substrates, and preparation methods, ranging between 10^4^ to 10^7^ Ω cm^2^. In comparison, suspended (black) planar lipid bilayers possess similar capacitance yet considerably higher resistance (10^6^ to 10^8^ Ω cm^2^) [54]. The lower resistance of SLBs can be understood as the result of inhomogeneities and defects in such large area membranes, which are considerably more physically stable compared to bilayers suspended in a liquid phase, hence can tolerate structural imperfections.

Although channel-free membranes are commonly characterized by simple resistive and capacitive elements, the electric response of experimental realizations of these membranes does not exactly follow this ideal model. The description of homogenous planar bilayers is complicated due to the complex microscopic nature of the lipid bilayer, its two interfaces with the surrounding electrolyte, and the submembrane electrolyte reservoir between the solid support and the lipid bilayer [21, 55]. Although the internal structure of membranes and the dielectric properties of each of its components can be modeled as an elaborate equivalent circuit, it is hard to unambiguously distinguish the contribution of each component to the EIS spectra [21, 56, 57]. Alternatively, the capacitive element describing the membrane can be generalized with a constant phase element (CPE) [21, 58, 59], with an impedance of the form 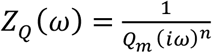. As *n* approaches the value of 1, *Z*_*Q*_ (*ω*) approximates an ideal capacitor. One should note that such a heuristic element lacks a concrete physical meaning. Moreover, they are, in principle, inadequate for the description of surface inhomogeneity and defects, for which more elaborate electrical network models are required [55, 60].

As shown in Fig. 1C, we model the electrical response of the system by assuming an overall impedance of the form *Z*(*ω*) = *Z*_*s*_ (*ω*) + *Z*_*m*_(*ω*), where *Z*_*s*_(*ω*) is the impedance associated with the substrate and electrolyte solution, and *Z*_*m*_ (*ω*) is the impedance associated with the lipid bilayer. The main contributors to *Z*_*s*_ (*ω*) are the supporting substrate sheet resistance *R*_*s*_, which includes a small contribution from the resistance of the electrolyte solution, and capacitance of the insulating oxide layer *C*_*s*_. Therefore, the impedance of the supporting substrate can be approximated to 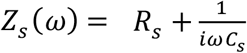. The impedance of the membrane is approximated by a resistive element *R*_*m*_ connected in parallel with a constant phase element 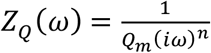, resulting in a total membrane impedance of the form 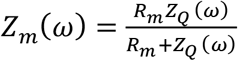. Fig. 2 shows the impedance spectra and the best fit of the equivalent circuit models, before and after bilayer formation, of two distinct representative preparations, differing mainly by their membrane resistance *R*_*m*_. The preparations of both the high *R*_*m*_ and low *R*_*m*_ membranes were identical, with the high *R*_*m*_ sample allowed for an additional stabilization time after membrane formation and before the staining procedure, as described in ‘Materials and methods’. As it will be shown, the electrical parameters describing the substrate and the membrane dictate the resulting magnitude and dynamics of the voltage drop on the membrane.

In the frequency domain, for an overall voltage drop *V*(*ω*) on the system, the voltage drop on the membrane would be:

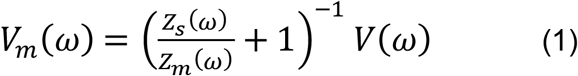

Assuming ideal resistive and capacitive components (*n* = 1), with 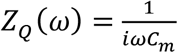, the voltage drop upon the substrate *V*_*s*_ and the membrane *V*_*m*_ follow simple dynamical equations:

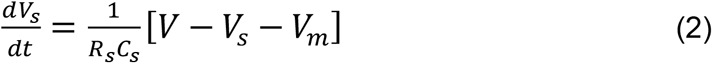

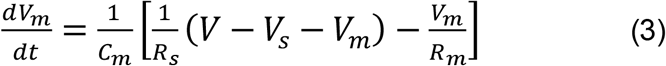

Where *V* is the time-dependent voltage drop on the whole circuit. Assuming the total voltage drop on the system is switched abruptly from 0 to *V*_0_ at *t* = 0, the step response of the potential drop on the membrane is

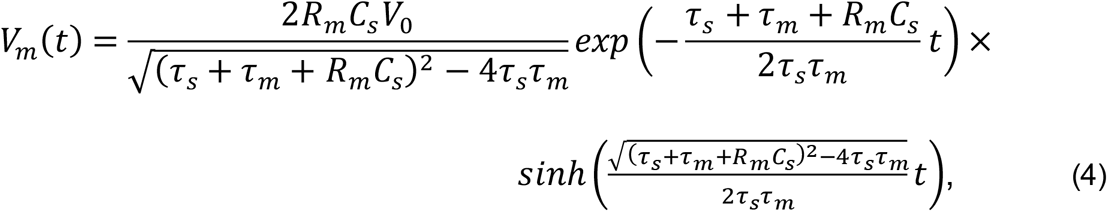

where *τ*_*m*_ = *R*_*m*_*C*_*m*_ and *τ*_*s*_ = *R*_*s*_ *C*_*s*_ are the time constant arising from the (parallel RC) membrane and (serial RC) substrate electrical properties, respectively. As it will be experimentally shown in the following section using optical measurements of the membrane potential, after a rapid initial rising period, the potential drop on the membrane decays exponentially as 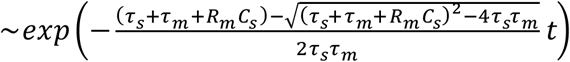. The decay of the membrane potential occurs due to the finite resistance of the membrane, which causes the voltage drop across the membrane to decrease, while the voltage drop across the insulating substrate increases and eventually approaches the value of the total voltage drop applied on the system. For the case of an infinitely resistive membrane, the potential drop on the membrane behaves as a simple charging capacitor 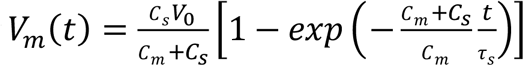, displaying no leakage of the membrane potential following its charging, with the total voltage drop shared between the membrane and the insulating substrate according to their capacitance values.

### Optical observation of membrane polarization

In order to demonstrate the ability to control the transmembrane voltage of the SLB, we used optical voltage-sensitive membrane dyes as an orthogonal measurement approach. Over the past five decades, a wide array of membrane potential indicators has been developed and applied for various biological membranes [61–64]. These optical indicators incorporate into the lipid bilayer, and display voltage-dependent optical properties, such as absorption and fluorescence. Here, we used VoltageFluor (VF) [65], a membrane dye that utilizes voltage-dependent photoinduced electron transfer (PET) between an electron donor and a fluorescent acceptor through a molecular wire (Fig. 3A). Specifically, we monitored the membrane potential of the electrically accessible supported membrane with VF2.1.Cl [65, 66], a VF derivative with a bi-exponential fluorescence decay curve that displays both intensity and lifetime changes upon changes in transmembrane potential. As a control, we used a similar derivative of the dye, VF2.0.Cl, which is spectrally similar to VF2.1.Cl, however, lacks the electron-rich donor, and therefore acts as a voltage-insensitive membrane dye [67]. Fig. 3 shows the fluorescence intensity and lifetime response of VF2.0.Cl and VF2.1.Cl upon electrical polarization of the supported membrane stained with 100 nM of either dye, using the high *R*_*m*_ membrane described in Fig. 2. In order to label only the leaflet closer to the bulk electrolyte, staining was performed externally for 5 min on a fully formed membrane, followed by ten rounds of buffer exchange in order to remove excess dye from the bath solution. All potentials were applied versus the Ag/AgCl electrode. Before each measurement, the detected region was photobleached, thus only the fluorescence of latterly diffusing dye molecules was measured. The absence of voltage-dependent changes in fluorescence of VF2.0.Cl implies that the voltage-dependent response of VF2.1.Cl can be attributed to changes in the directional transmembrane potential, and not to other artifacts, such as local change of ion concentrations upon polarization of the working electrode. The change in fluorescence intensity of VF2.1.Cl upon a total applied polarization of 500 mV was measured to be Δ*F*/*F* ≈ 47 ± 1.3 %, averaged over 30 polarization cycles at three regions of the supported membrane. The fluorescence amplitude-averaged lifetime showed an increase from 0.64 ± 0.03 ns at –250 mV to 0.70 ± 0.02 ns at 250 mV, corresponding to a relative change of Δ*τ*/*τ* ≈ 10 ± 4.1% upon a total applied polarization of 500 mV.

**Fig. 3.**
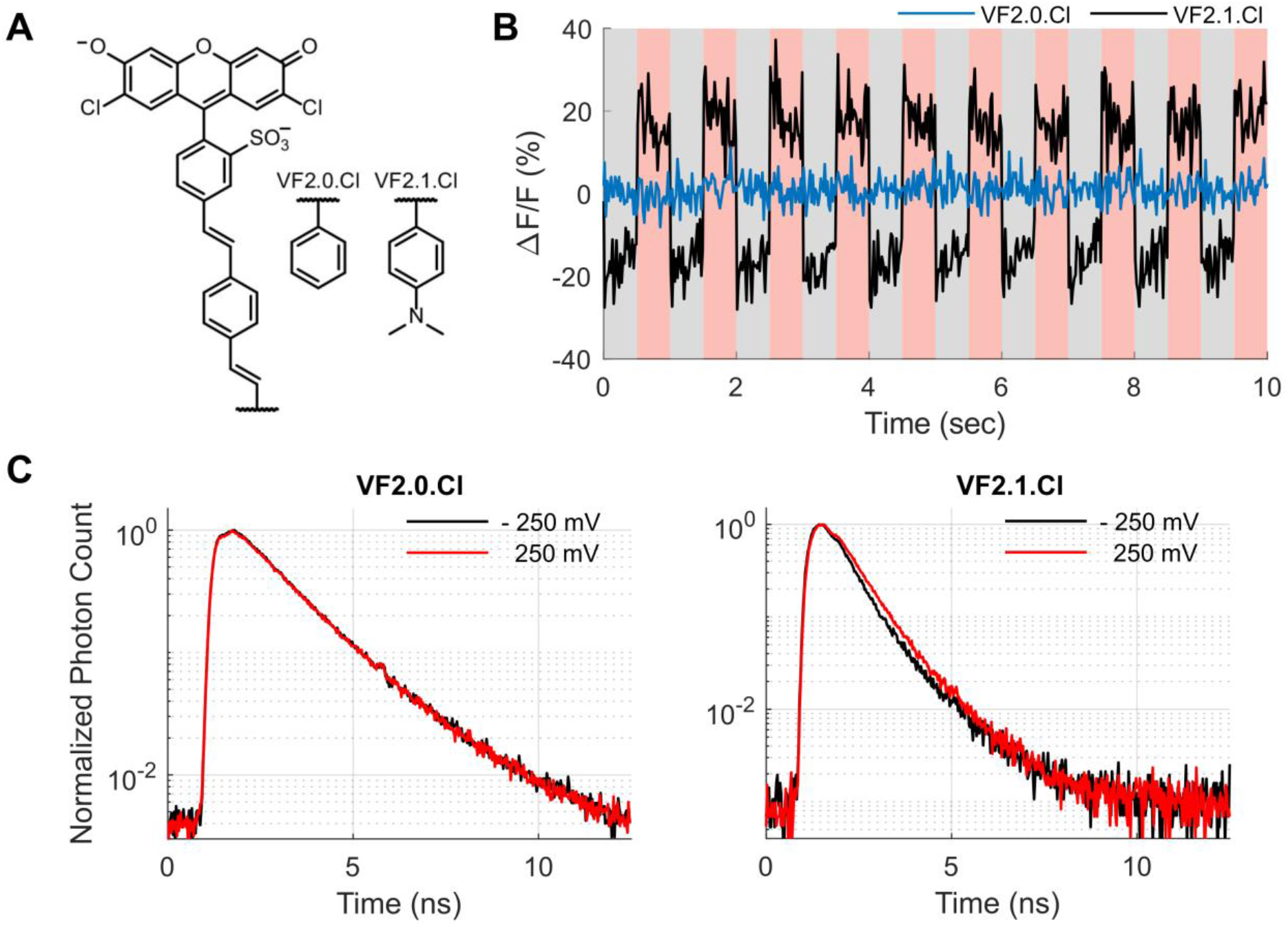
Optical monitoring of changes in membrane potential. **(A)** Structure of the two VF membrane dyes used in this study. Both dyes are composed of a fluorophore conjugated to a molecular wire, with (VF2.1.Cl) or without (VF2.0.Cl) an electron-rich aniline donor, rendering the dyes to be either voltage sensitive or insensitive, respectively. **(B)** The fluorescence emission intensity of a voltage-sensitive (VF2.1.Cl, black) and a voltage-insensitive (VF2.0.Cl, blue) membrane dyes upon electrical polarization of the stained membrane. The total voltage drop upon the electrochemical system was alternated between −250 mV (gray-shaded area) and +250 mV (red-shaded area). **(C)** The fluorescence lifetime decay curves of both dyes at a voltage drop of −250 mV (black) and 250 mV (red), averaged over 30 polarization cycles, showing lifetime response of VF2.1.Cl to changes in membrane potential.

As discussed in the previous section, one should note that the voltage drop across the membrane (the transmembrane potential) is necessarily smaller than the overall applied voltage, since the overall voltage drop is distributed among the different electrical components constituting the electrochemical system. Fig. 4 shows the averaged fluorescence intensity response of VF2.1.Cl of both the high and low *R*_*m*_ membranes characterized in Fig. 2. As discussed in the previous section, both the magnitude and dynamics of the voltage drop on the membrane are highly dependent on the electrical properties of the metal-insulator-bilayer system. In particular, the membrane resistance *R*_*s*_, which significantly varies between preparation methods of the lipid bilayer, dictates to a great extent the leakage of membrane potential to the substrate over time. The substrate capacitance *C*_*s*_, which is controlled by the fabrication process, has to be as large as possible in order to increase the actual voltage drop on the membrane, as demonstrated in Fig. S4. The predicted membrane potential magnitude and dynamics can be calculated using Eqs. 1 and 4, using the fitted parameters of the equivalent circuit. Since the CPE exponent *n* is close to 1, as shown in Fig. 2, the more complex CPE description is approximated well by an ideal capacitive element. Hence, as shown in Fig. 4, the simplified model, in which the membrane is modeled by ideal resistive and capacitive components, is in good agreement with the optically observed membrane electric polarization.

**Fig. 4.**
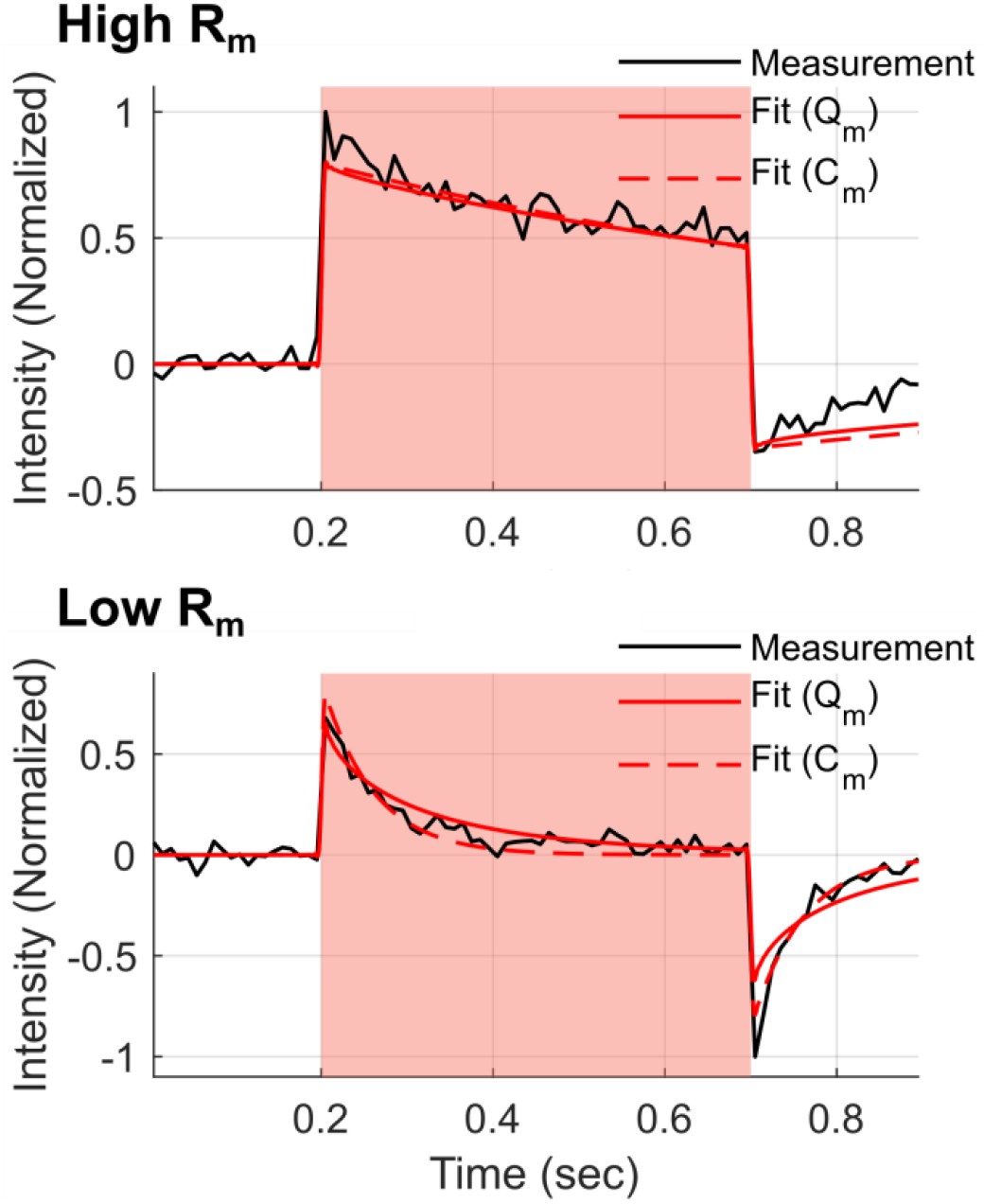
Optical monitoring of membrane potential dynamics. The averaged fluorescence emission intensity (black) change upon polarization to +500 mV (red-shaded area) of the high (upper panel) and low (lower panel) resistance membranes shown in Fig. 2, averaged over 20 polarization cycles. The expected response to such polarization pulse is calculated by solving the dynamical equations of the equivalent circuit model of the membrane, using the best fit parameters described in Fig. 2, assuming the change in transmembrane potential follows the voltage drop on the membrane *V*_*m*_(*t*). The behavior of the membrane potential *V*_*m*_(*t*) upon electric polarization is shown for a membrane modeled both with a constant phase element *Q*_*m*_ (solid red) and an ideal capacitive element *C*_*m*_ (dashed red).

For the high *R*_*m*_ membrane demonstrated in Figs. 2 and 3, assuming the membrane potential step response described in Eq. 4, a total voltage drop of 500 mV results in membrane polarization of approximately 300 mV at its peak. Even after accounting for the smaller membrane voltage drop due to circuit dynamics, the measured optical response to changes in membrane potential is smaller than previously published results from patch-clamped cells stained with VF2.1.Cl. In HEK293T cells, the change in fluorescence intensity of VF2.1.Cl was measured to be Δ*F*/*F* ≈ 27 % per 100 mV, and the fluorescence lifetime showed a sensitivity of 3.5 ± 0.08 ps/mV and a lifetime of 1.77 ± 0.02 ns at 0 mV, corresponding to a relative change of Δ*τ*/*τ* ≈ 22.4 ± 0.4% per 100 mV [65, 66]. This discrepancy can be attributed to several factors. Since the interface and environment on both sides of the supported membrane are highly asymmetrical, assigning one value for the transmembrane potential may be an oversimplification. For example, one could test and compare the optical response of voltage-sensitive membrane dyes in each of the bilayer leaflets, or increase the electrolyte reservoir underneath the membrane by distancing the bilayer from the solid support [19]. Another important consideration should be the effect of the conducting layer on the fluorescence of the optical probe. In particular, metal-induced energy transfer (MIET) [44, 68] increases the radiative transition rate of the fluorophore as the fluorophore gets closer to the conducting layer, resulting in a shorter lifetime. Fig. 5A shows the lifetime of VF2.0.Cl and VF2.1.Cl in membranes supported on glass and the Ti/Ta_2_O_5_ substrate used for the measurements described above. The lifetime of the mono-exponential VF2.0.Cl reduced from 2.40 ± 0.08 ns to 1.12 ± 0.07 ns for membranes supported on glass and the Ti/Ta_2_O_5_ substrate, respectively. The bi-exponential decay curve VF2.1.Cl, which has an amplitude-averaged lifetime of 0.78 ± 0.04 ns for membranes supported on glass, reduced to 0.48 ± 0.003 ns when supported by the Ti/Ta_2_O_5_ substrate. However, the lifetime of VF2.1.Cl on the same substrate was measured to be 0.67 ± 0.02 ns once the Ti/Ta_2_O_5_ substrate was electrically connected to the potentiostat and the total voltage drop on the system was set to 0 mV. A possible explanation for the difference of the fluorescence decay between the two measurements is the native surface potential of the electrode-membrane system, which is different from the case where the system is electrically set to a constant overall potential of 0 mV.

**Fig. 5.**
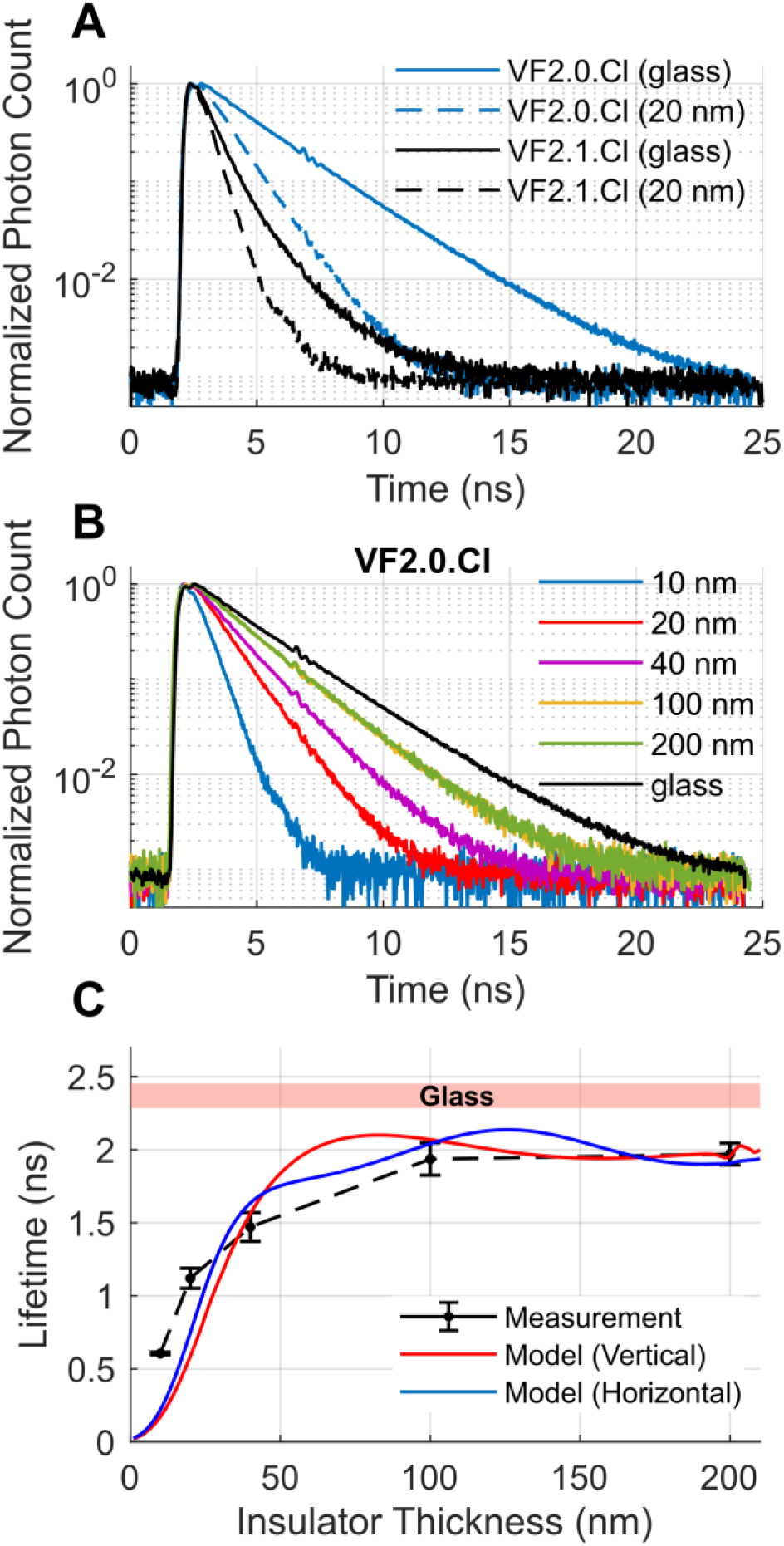
Metal-induced energy transfer (MIET) influence on the fluorescence lifetime decay. **(A)** The lifetime decay curves of VF2.0.Cl (blue) and VF2.1.Cl (black), staining a membrane supported by a glass (solid line) or Ti/Ta_2_O_5_ (dashed line) substrate, with 20 nm insulating layer of Ta_2_O_5_. **(B)** The lifetime decay curves of VF2.0.Cl supported by Ti/Ta_2_O_5_ substrates with varying thicknesses of the insulating layer Ta_2_O_5_. The lifetime decay constant for each insulating layer thickness, averaged over five different regions of the membrane, are shown in **(C)** (black), with the red-shaded area indicating the lifetime distribution of VF2.0.Cl in a membrane supported on bare glass. The theoretically calculated dependence of the fluorescence lifetime on the insulator thickness is shown for the case of an emission dipole that is orthogonal (red) and parallel (blue) to the substrate, assuming a quantum yield of 0.56 and a free space lifetime of 3.29 ns.

The dependence of the fluorescence lifetime on the distance between the membrane dye and the conducting Ti layer was theoretically modeled for VF2.0.Cl, which has a simpler mono-exponential decay curve. Fig. 5B shows the fluorescence decay curve of VF2.0.Cl in membrane supported on a Ti/Ta_2_O_5_ substrate with varying thicknesses of the insulating layer Ta_2_O_5_, displaying the shortening of the fluorescence lifetime due to MIET as the membrane dye is closer to the conductive layer. Fig. 5C shows the fitted fluorescence decay constant for each decay curve, along with the theoretically calculated lifetime for a model emission dipole [69, 70]. We note that the discrepancy between the measurement and the theoretical model cannot be explained merely as the result of inaccurate knowledge of emission dipole orientation, free space fluorescence lifetime, or quantum yield, as was checked by altering these fluorophore-dependent parameters in the model calculations. A possible explanation could be that the assumed imaginary part of the refractive index of the conducting layer is inaccurate, which would greatly change the dependence of the fluorescence lifetime on the distance to the conducting layer. Here, we chose this parameter using the Brendel–Bormann model [71]. However, as shown in Fig. S3A, the ~8 nm Ti layer displays a non-uniform electron density, most likely due to the consecutive Ta_2_O_5_ deposition. This non-homogeneity may cause the optical properties of the Ti layer to differ from the ones assumed in the model, and therefore alter the effect of MIET on the fluorescence lifetime.

The lifetime dependence of the fluorescent probe on its distance from the conducting layer can be utilized to measure its axial location with sub-nm resolution [72], as will be discussed in the following section. In addition, one should take this change in radiative transition rate into account when considering quantitative optical measurements based on fluorescent quenching like Förster resonance energy transfer (FRET) or PET. The exact nature of this effect will depend upon the complete description of electronic transitions from the photoexcited state. For example, we consider a simple model in which a photoexcited state relaxes either radiatively or non-radiatively with a total de-excitation transition rate of *k*_*T*_ = *k*_*R*_(*z*) + *k*_*NR*_ + *k*_*ET*_ (*V*), where*k*_*R*_(*z*) is the radiative rate which depends upon the distance of the fluorophore from the conducting surface, *k*_*NR*_ is the non-radiative relaxation rate, and *k*_*ET*_ (*V*) is a voltage-dependent electron transfer rate to a charged-separated state, assuming the charge-separated state relaxes non-radiatively [73]. The average emission intensity *F* of such fluorophore is proportional to its quantum yield, i.e. *k*_*R*_(*z*)/*k*_*T*_. The relative change in fluorescent emission due to small changes in the applied voltage is Δ*F*/*F* = −Δ*k*_*ET*_ (*V*)/*k*_*T*_, where Δ*k*_*ET*_ (*V*) is the corresponding change in the electron transfer transition rate *k*_*ET*_ (*V*) as a function of applied voltage. Therefore, an increased radiative transition rate, such as in the case of MIET, inevitably reduces the sensitivity of such optical probes to changes in membrane potential.

### Future work – optical investigation of voltage-dependent protein dynamics and biologically relevant model membranes

Fluorescence microscopy and spectroscopy are frequently used in the study of membranes and their associated proteins. In particular, FRET provides a powerful approach for nanoscale measurements of molecular structure and dynamics of biomolecules such as DNA, RNA, proteins, and their interaction [74–78]. While FRET allows for the measurement of the distance between donor and acceptor molecules, MIET allows for nanoscale axial localization of fluorescent probes above the conducting substrate [43, 44]. In particular, using graphene as the conducting substrate results in significant dependence of the fluorescence lifetime to sub-nm changes in the axial direction [72, 79], making this method highly relevant for the study of membrane proteins. Applying optical techniques such as FRET and MIET utilizing the platform described in this work can allow for novel experiments that probe voltage-induced molecular dynamics of membrane proteins. Beyond voltage-gated ion channels, this experimental approach could investigate the role of membrane potential and electrostatics in the conformation of membrane proteins [38–41]. In addition to membrane and protein biophysics, combining optical and electrochemical tools can be used for studying the physicochemical properties of solid-electrolyte interfaces. Such studies were performed using electrochemiluminescence [80, 81] and electric-double-layer-modulation [82] microscopies, as well as many other optical techniques [83]. In addition to fluorescence microscopy of fluorescent probes that are inherently environment-sensitive, applying ultrasensitive methods such as FRET and MIET with probes that are electrically responsive due to their net charge can allow for the study of interfaces at a sub-nm resolution.

Although synthetic lipid bilayers, such as the ones prepared in this work, provide a clean and robust experimental system, they fail to mimic important features of real biological membranes, which contain a rich complexity of biological membranes such as different phospholipid headgroups, saturation levels of their tails, membrane proteins, cholesterol, and carbohydrates [84]. Such realistic membranes will include lateral inhomogeneities and macroscopic and nanoscopic domains of different membrane fluidity and phase. More biologically relevant membranes can be achieved by either additional synthetic [85] or biologically-derived components [86], or by extracting the components from real cell membranes by induced extracellular vesicles [87]. One should note that the addition of foreign components to the bilayer film, such as labeled fluorescent probes, can affect the overall functional properties of the membrane [88]. Hence, low probe concentrations and proper control experiments should be considered when adding extrinsic fluorescent probes. In addition, enhanced membrane stability and flexibility can be achieved through the introduction of a polymer cushion layer or a tethering layer that anchors one leaflet to the substrate [16, 17, 89–91]. Similar to the work shown here, these tethered membranes can also be electrochemically characterized when deposited on a conductive substrate [55, 92–94].

## Conclusion

Combining both electrochemical and optical tools for studies that utilize SLBs offer novel possibilities for voltage-dependent investigations of membranes and their associated biomolecules. While electrochemical tools allow for electrical characterization and control of the membrane and the supporting substrate, optical measurements offer an orthogonal approach for probing membranes, membrane proteins, and their dependence on external signals. In this work, we established an electrochemical-optical platform for SLBs and demonstrated electrical control of the membrane potential by independently monitoring it with voltage sensitive dyes. We showed that the membrane potential is governed by the circuit dynamics of the electrochemical system, which can be understood by its characterization via EIS. In addition, we demonstrated that the conducting electrode layer affects the fluorescent emission of the optical probe through MIET, and discussed possible resulting implications. Such electrically and optically addressable membranes can provide a platform for a vast array of ensemble and single-molecule optical methods for the study of folding and interaction of membrane proteins.

## Contributions

S.Y. prepared the samples and performed optical and electrochemical experiments.

A.M. prepared the vesicles. J.K. fabricated the metal-insulator electrodes.

E.T. performed AFM for surface characterization. T.C. and J.E. analyzed the MIET data.

E.W.M. synthesized the membrane dyes. S.Y. and S.W. designed and planned the study. All co-authors participated in writing the manuscript.

## Acknowledgements

We thank Dr. Maria Tkachev and Dr. Ilana Perelshtein for performing cross-sectional HR-TEM imaging of the electrode and Dr. Ayelet Atkins for performing cryo-EM imaging of the vesicles. This work has received funding from the European Research Council (ERC) under the European Union’s Horizon 2020 research and innovation program under grant agreement No. 669941, by the Human Frontier Science Program (HFSP) research grant RGP0061/2015, by the BER program of the Department of Energy Office of Science grant DE-SC0020338, by the STROBE National Science Foundation Science & Technology Center, Grant No. DMR-1548924, by the Israel Science Foundation (ISF) Grant 813/19, and by the Bar-Ilan Research & Development Co, the Israel Innovation Authority, Grant No. 63392.

## Data availability

Raw data of all EIS, imaging and time-resolved fluorescence measurements presented in this work is available at Zenodo: 10.5281/zenodo.5769883. Home-written code (MATLAB) for the analysis shown in this work is available upon request.

