## Supplementary Material for "Electrically Controlling and Optically Observing the Membrane Potential of Supported Lipid Bilayers"

### Supplementary Figures

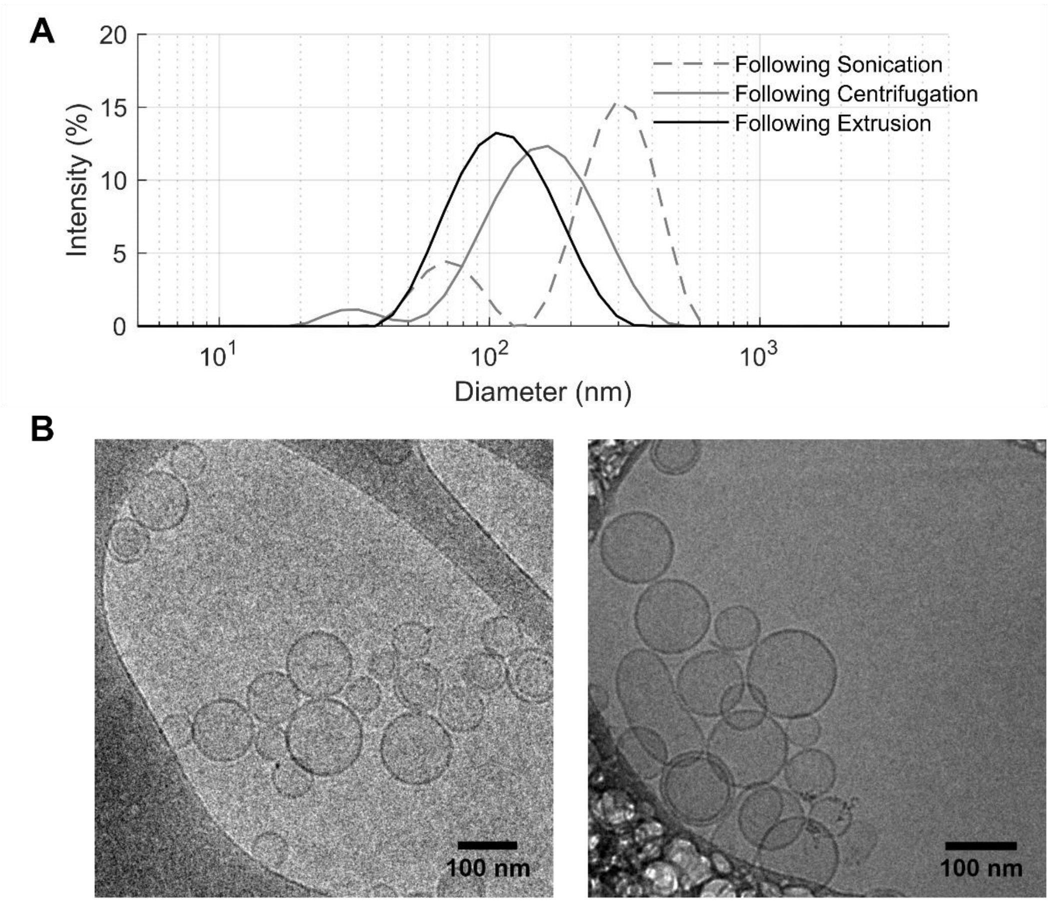


**Fig. S1.** Size characterization of the vesicles used for the formation of supported lipid bilayers. **(A)** Dynamic light scattering intensity distribution of the vesicles after each preparation step, as described in 'Materials and methods'. Vesicles following extrusion were used for the formation of supported lipid bilayers. In order to ensure repeatability between samples, the total scattered intensity was used as an indicator for the vesicle concentration. The size populations (and the polydispersity index, PDI) after each step are, following sonication - two populations at 311 ± 85 nm and 71 ± 16 nm (PDI: 0.408), following centrifugation - two populations at 171 ± 72 nm and 33 ± 8 nm (PDI: 0.216), and following extrusion - one population at 121 ± 50 nm (PDI: 0.155). The refractive indices used for calculating size distributions were 1.45 and 1.331 for the vesicles and the buffer, respectively. **(B)** Cryogenic transmission electron microscopy images of the extruded vesicles.

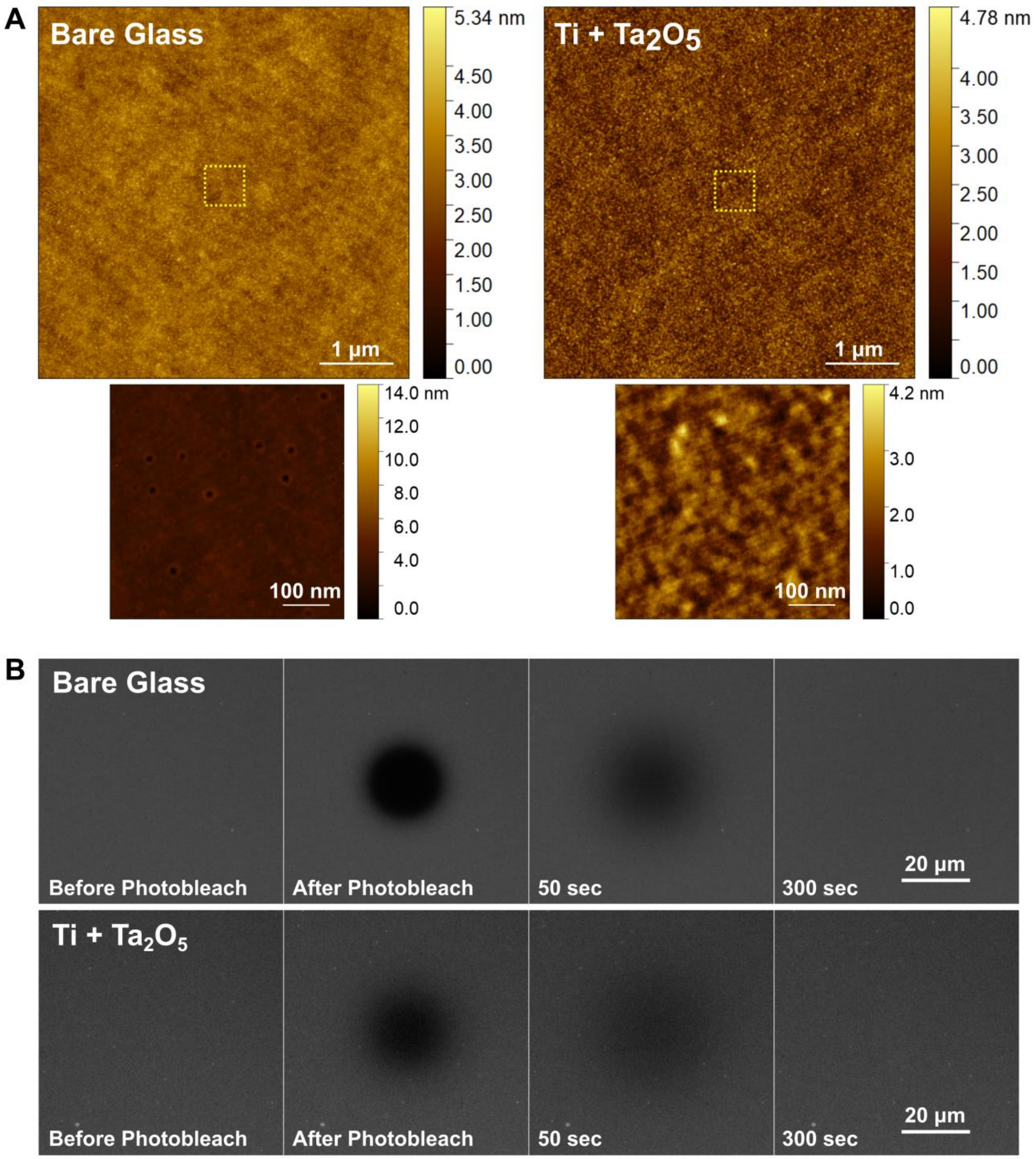


**Fig. S2.** Surface characterization of the membrane-supporting metal-insulator substrate. **(A)** Atomic force microscope (AFM) images of the substrate before (left) and after (right) the deposition of ~8 nm Ti and ~20 nm $Ta_2O_5$. Zoomed-in scans of the area indicated by a yellow dashed square are shown on the bottom panel. The RMS roughness, averaged over three scans of a 5 x 5 µm$^2$ area, is 0.38 ± 0.01 nm for bare glass, and 0.47 ± 0.01

nm for the Ti/$Ta_2O_5$ substrate. The bare glass substrate shows homogenous holes throughout the substrate which are likely generated during the processing of the glass. These holes are covered by the deposited layers and are not seen in the final Ti/$Ta_2O_5$ substrate. **(B)** Fluorescence recovery after photobleaching (FRAP) of POPC:POPS lipid bilayers stained with 1 µM of rhodamine-labeled lipids (18:1 Liss Rhod PE; Avanti Polar Lipids, Alabaster, AL), showing the fluidic phase of the supported membranes. Staining was performed externally on fully-formed membranes in order to label solely the leaflet facing the electrolyte, as was done for VF2.0.Cl and VF2.1.Cl in this work. Bleaching of the membrane supported by the Ti/$Ta_2O_5$ substrate was partial due to the fact that the excitation intensity had to be limited, as high intensity excitation caused a physical disruption of the membrane, likely due to metal-enhanced local heating of the membrane.

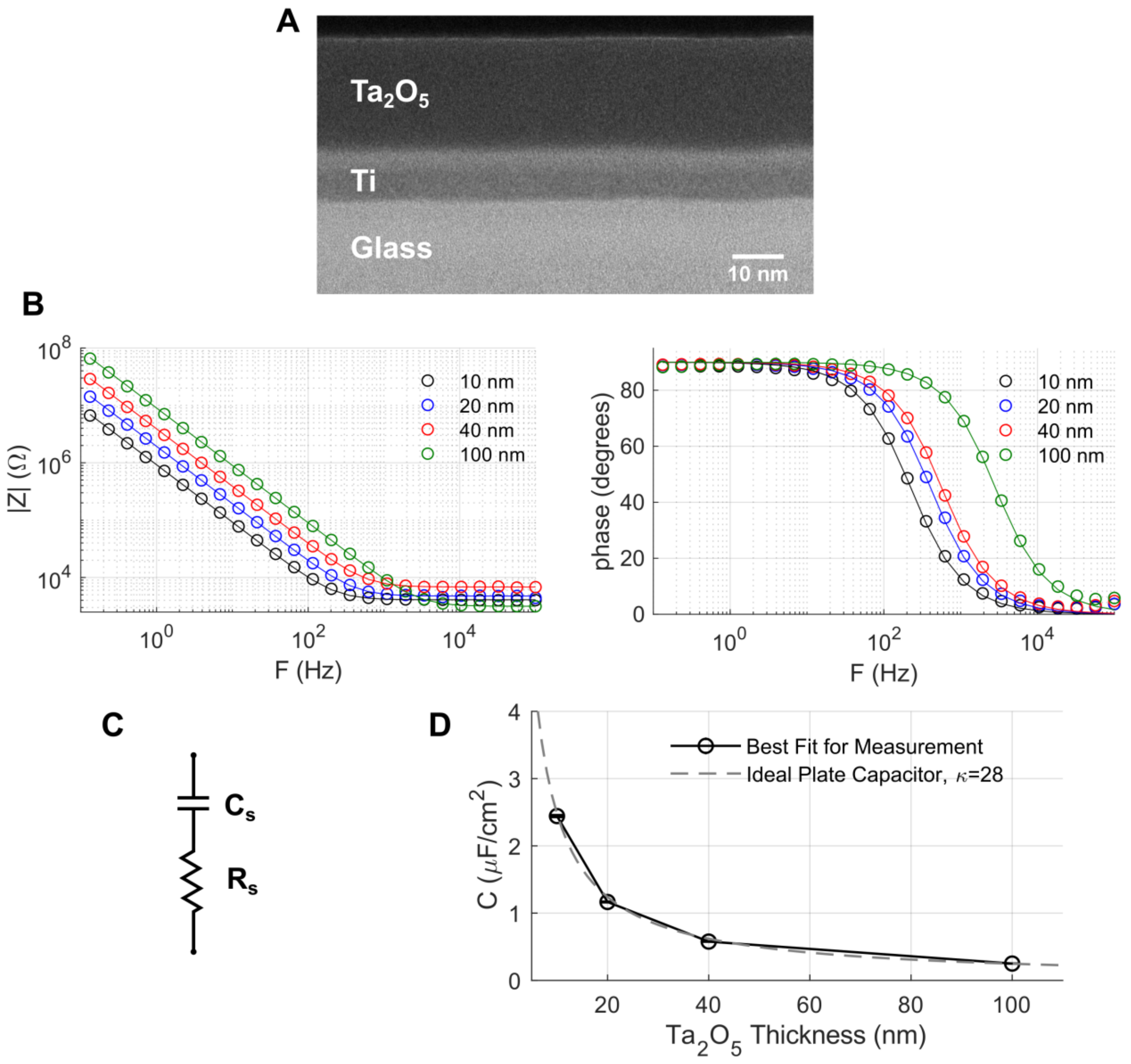


**Fig. S3.** Electrical characterization of the membrane-supporting metal-insulator substrate. **(A)** Cross-sectional transmission electron microscope (TEM) image of a lamella from the electrode (~8 nm Ti and ~20 nm $Ta_2O_5$) milled using gallium focused ion beam (FIB). **(B)** Electrochemical impedance spectroscopy (EIS) measurements of the Ti/$Ta_2O_5$ electrode with varying thicknesses of the insulating $Ta_2O_5$ layer (open circles). The left and right panels show the impedance amplitude and phase, respectively. The solid lines represent the impedance spectra of the best fitted model using the model equivalent circuit shown in **(C).** All measurements were performed in 10 mM Tris with 100 mM NaCl (pH 8). The best fit capacitance of the insulating layer $C_s$ as a function of it the layer thickness is shown in **(D)**, with the dashed line representing the capacitance per unit area of an ideal plate capacitor, $C = \kappa\varepsilon_0/d$, where $\kappa$ is the dielectric constant, $\varepsilon_0$ is the vacuum permittivity, and $d$ is the thickness of the layer. The fitted capacitance values show good agreement with an ideal plate capacitor model with a dielectric constant of $\kappa = 28$, which is a typical value for $Ta_2O_5$.

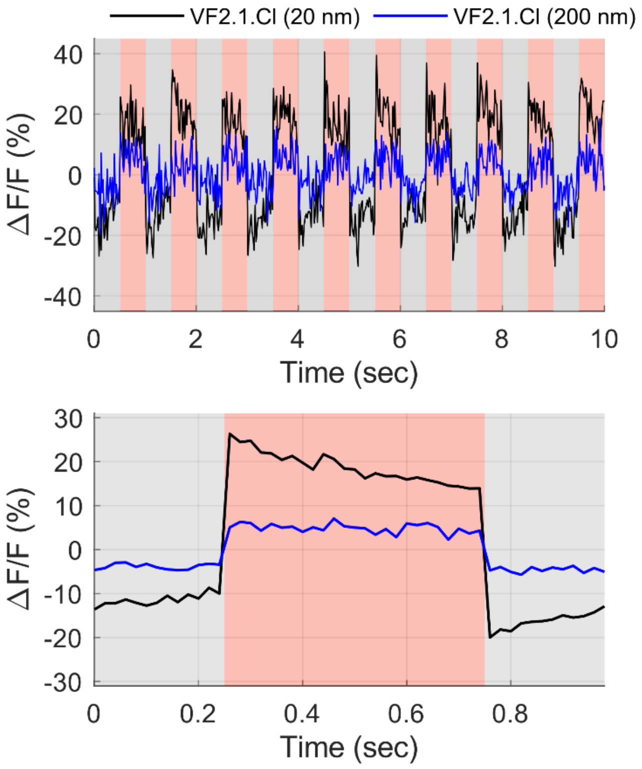


**Fig. S4.** The dependence of the supported membrane's voltage drop on the insulating layer. The top panel shows the fluorescence emission intensity of VF2.1.Cl upon electrical polarization of a stained membrane for both a 20 nm (black) and 200 nm (blue) insulating $Ta_2O_5$ layer, with the average change shown in the bottom panel (averaged over 30 polarization cycles). The total voltage drop upon the electrochemical system was alternated between -250 mV (gray-shaded area) and +250 mV (red-shaded area). The averaged response for the total 500 mV polarization is $\Delta F/F \approx 46\%$ for the 20 nm insulating layer and $\Delta F/F \approx 11\%$ for the 200 nm insulating layer. EIS measurements of both samples before and after membrane deposition were fitted to an equivalent circuit as shown in Fig. 2. Best fit parameters for the case of an ideal capacitive membrane were $R_s = 249.4\ \Omega\, cm^2,\ C_s = 1.2\ \mu\text{F}\, cm^{-2}$, $R_m = 457.9\ k\Omega\, cm^2,$ $C_m = 0.79\ \mu\text{F}\, cm^{-2}$ for the 20 nm insulating layer, and $R_s = 218.8\ k\Omega\, cm^2,\ C_s =$ $0.12\ \mu\text{F}\, cm^{-2}$, $R_m = 131\ k\Omega\, cm^2,\ C_m = 0.96\ \mu\text{F}\, cm^{-2}$ for the 200 nm insulating layer. Assuming these values for the equivalent electrical circuit, and using Eq. 4, the peak voltage drop on the membrane supported by the 20 nm insulating layer is expected to be approximately ~5.5 higher than for the membrane supported by the 200 nm insulating layer.
